# Flexible, multi-use, PCR-based nucleic acid integrity assays based on the ubiquitin C gene

**DOI:** 10.1101/168195

**Authors:** Mario Van Poucke, Luc J. Peelman

## Abstract

Nucleic acid integrity assessment is an important aspect of quality control for many applications in molecular biology. A number of methods exist (electrophoresis- or PCR-based), but they are not universally applicable. Some of them need huge amounts of input, process certain amount of samples, require expensive equipment, only work specifically on (c)DNA, (m)RNA, certain species or certain tissues, or produce fragments covering a small length range. We investigated if the ubiquitin C gene (UBC) could be used to develop flexible, multi-use PCR-based (deoxy)ribonucleic acid integrity assays. UBC gene analysis (in human, mouse, pig, cow, horse, sheep, dog and cat) shows that UBC is a highly conserved and ubiquitously expressed gene (reference gene in RT-qPCR), that encodes a polyubiquitin precursor (containing tandem repeats of at least 5 ubiquitin monomers of 228 bp) in a single exon. On average, ubiquitin monomers show a nucleic acid sequence identity of 96% at intraspecies level and 93% at interspecies level. Based on a multiple alignment of all monomer ubiquitin sequences of all investigated species, we could design a single degenerated primer pair generating PCR amplicons of 137, 365, 593 and 821 bp on low amounts of high quality DNA of all investigated species (down to 10 pg) and on cDNA reverse transcribed from high quality RNA from different tissues (e.g. heart, liver, brain, kidney). Increasing levels of nucleic acid degradation resulted in a decrease of amplification products starting with the longer amplicons. We conclude that UBC is suited to develop a single, cheap, universal assay to estimate the presence, integrity and amplificability of native mammalian nucleic acids. In addition, we used the same strategy to design a similar assay to check the quality of bisulfite treated mammalian DNA.

## INTRODUCTION

Nucleic acid integrity assessment is an important aspect of quality control faced by every researcher in the decision if a sample is suitable for many of the, sometimes expensive and/or time-consuming, applications in molecular biology. Existing integrity assays rely on electrophoresis or PCR, but they are not universally applicable/available [1–3]. Gel electrophoresis is cheap and easy to perform, but requires high amounts of sample. Chip electrophoresis is more sensitive and needs only a limited amount of sample, but is expensive and requires specialized equipment. In contrast with PCR-based assays, they do not check for PCR amplificability. Traditional PCR assays are cheap, easy to perform, sensitive and require only a limited amount of sample, while qPCR or dPCR assays might be more precise, but more complex, more expensive and require specialized equipment. Because published PCR-based assays are developed application-dependent, they only work specifically on DNA, cDNA, certain species, certain tissues or produce fragments covering a small length range [1–3]. As a result, until now, laboratories that want to check PCR amplificability have to implement a battery of different integrity assays to cover all their experiments.

The ubiquitin C gene (UBC) is a highly conserved and ubiquitously expressed gene (frequently used as a reference gene in RT-qPCR), that encodes a polyubiquitin precursor in a single exon [4–5]. Here, we want to investigate if the UBC gene could be used to develop a single, cheap, multi-use PCR-based assay to estimate the presence, integrity and amplificability of DNA (isolated from man, mouse, pig, cow, horse, sheep, dog or cat) and cDNA (reverse transcribed from RNA isolated from any tissue type and reflecting RNA integrity).

## MATERIALS AND METHODS

### DNA isolation

DNA was isolated from blood by performing a proteinase K digest followed by a phenol/chloroform extraction and an ethanol precipitation. DNA was artificially degraded with 0,1 U RQ1 DNase (Promega). DNA used for bisulfite treatment was isolated with the Quick-DNA Miniprep Plus kit (ZymoResearch). Bisulfite treatment was performed with the EZ DNA Methylation-Lightning Kit (ZymoResearch). Purity and concentration was measured with Nanodrop (Isogen). Integrity was estimated by analysing 1 μg of DNA on a 1% agarose gel.

### RNA isolation and cDNA synthesis

Total RNA was isolated using the Aurum Total RNA Fatty and Fibrous Tissue Kit (Bio-Rad), including an on-column DNase treatment to remove genomic DNA. Purity and concentration was measured with Nanodrop (Isogen). Integrity was estimated by analysing 1 μg of RNA on a 1% agarose gel. One μg of total RNA was converted to cDNA with the ImProm-II cDNA synthesis kit (Promega), using both oligo dT and random primers. cDNA was 10 times diluted with water.

### PCR

UBC gene sequences were retrieved via NCBI Gene [6]. Pairwise sequence alignment was performed with NCBI Blast [7] and multiple sequence alignment with ClustalW [8]. Primers based on native DNA were designed with Primer-Blast [9] and primers based on bisulfite treated DNA with BiSearch [10]. PCR assays were optimised and performed with TEMPase Hot Start DNA Polymerase (VWR) on a S1000 Thermal Cycler (Bio-Rad). Assay results were validated with primers amplifying other genes. All assay details are listed in Additional file 1.

### Agarose gel electrophoresis

DNA, RNA and PCR products were all analysed by 1% agarose gel electrophoresis.

## RESULTS AND DISCUSSION

### UBC gene *in silico* analysis

Dot matrix views of intraspecies pairwise nucleic acid sequence alignment of human, murine, porcine, bovine, equine, ovine, canine and feline UBC gene sequences show that all investigated UBC genes encode at least 5 tandem repeats of the UBC monomer of 228 bp (Additional file 2). Intraspecies pairwise nucleic acid sequence alignments of the individual UBC monomer sequences show that they are highly conserved at intraspecies level, with 96% nucleic acid sequence identity averaged over the investigated species (except for the 10^th^ UBC monomer in mouse, that was left out for further analysis). Interspecies pairwise nucleic acid sequence alignments of the individual UBC monomer sequences show that, with an average of 93% nucleic acid sequence identity, they are even highly conserved at interspecies level. Multiple nucleic acid sequence alignment of all individual UBC monomer sequences of the investigated mammals shows positional conservation (Additional file 3).

### UBC integrity assay design

The rationale behind the assay design is to choose primers that are able to bind to all UBC monomers of all investigated species in order to amplify a series of fragments of different lengths due to the presence of the tandem repeats (Figure 1). Since these tandem repeats contain uninterrupted coding sequences and UBC is ubiquitously expressed, primers can amplify both DNA and cDNA reverse transcribed from RNA isolated from any tissue (Additional file 4).

**Figure 1.**
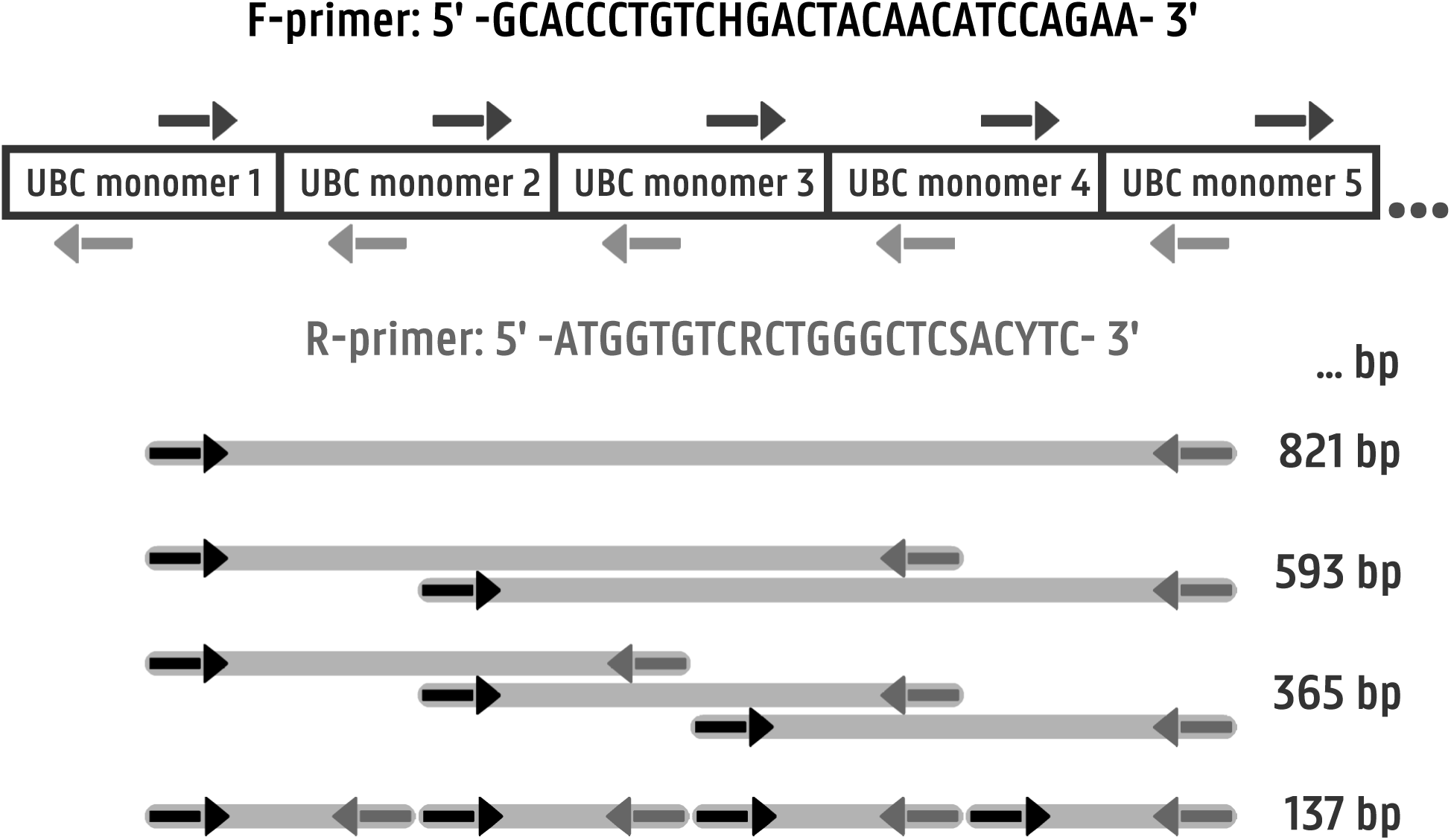
UBC integrity assay design strategy, showing primer sequences and amplicon lengths.

The amplification rate of the different fragments will be related to the quality of the template. High quality nucleic acids will generate a ladder of amplicons. Decreasing quality will result in a decrease of both the number and the concentration of the amplicons, starting with the longer fragments. By analysing the amplification pattern of this single assay, it is possible to check the presence, integrity and amplificability of the template, being DNA or cDNA from any tissue of any of the investigated mammals.

Because of the presence of single ubiquitin monomer sequences in several other genes, primers were chosen not to be able to amplify single monomers in order not to implicate the amplification of the longer UBC fragments. The human UBC nucleic acid sequence containing the first two monomers was used to design primers. The forward primer was forced to be chosen in the last 114 bp of the first monomer and the reverse primer in the first 114 bp of the second monomer, in order to allow only amplification of ubiquitin tandem repeats (Figure 1). Potential primers were manually checked for variation based on the multiple nucleic acid sequence alignment of all individual UBC monomer sequences of the investigated mammals (Additional file 3). Optimal primers should contain no or only a few variations, preferably at their 5’-end, and they should not be present in 5 consecutive monomers in order not to implicate amplification of fragments up to 1000 bp. Variation outside these 5 consecutive monomers, in species with more than 5 tandem repeats, was ignored. Finally, a primer pair in regions with the least variation was ordered, with degenerate nucleotides where variation occurred, aiming to generate amplicons of 137, 365, 593 and 821 bp (Figure 1; Additional file 1.A).

A similar strategy was followed to design the UBC bisulfite integrity assay, aiming to generate amplicons of 105, 333, 561 and 789 bp on the bisulfite treated UBC DNA sense strand (Additional file 1.A).

### Validation of the UBC integrity assay

Gradient PCR showed that all desired amplicons were generated on 100 ng high quality porcine DNA using a wide annealing temperature (Ta) range (68°C was chosen as optimal Ta; Additional File 5.A). The optimized assay was used to check its sensitivity by performing the assay on a 1/10 serial dilution series from 100 ng down to 100 fg of high quality porcine DNA. Agarose gel analysis shows that all desired amplicons were generated down to 10 pg of template (equivalent to DNA from 2 cells). One pg input resulted in the amplification of only the smaller bands, while 100 fg resulted in no amplification products (Figure 2.A). Species specificity was confirmed with 100 ng of high quality DNA isolated from all the investigated mammals as a template (Figure 2.B) and tissue specificity was confirmed with cDNA (equivalent to 10 ng RNA) from jejunum, heart, mucosa, liver, brain, kidney, neutrophil and lymph node (Figure 2.D).

**Figure 2.**
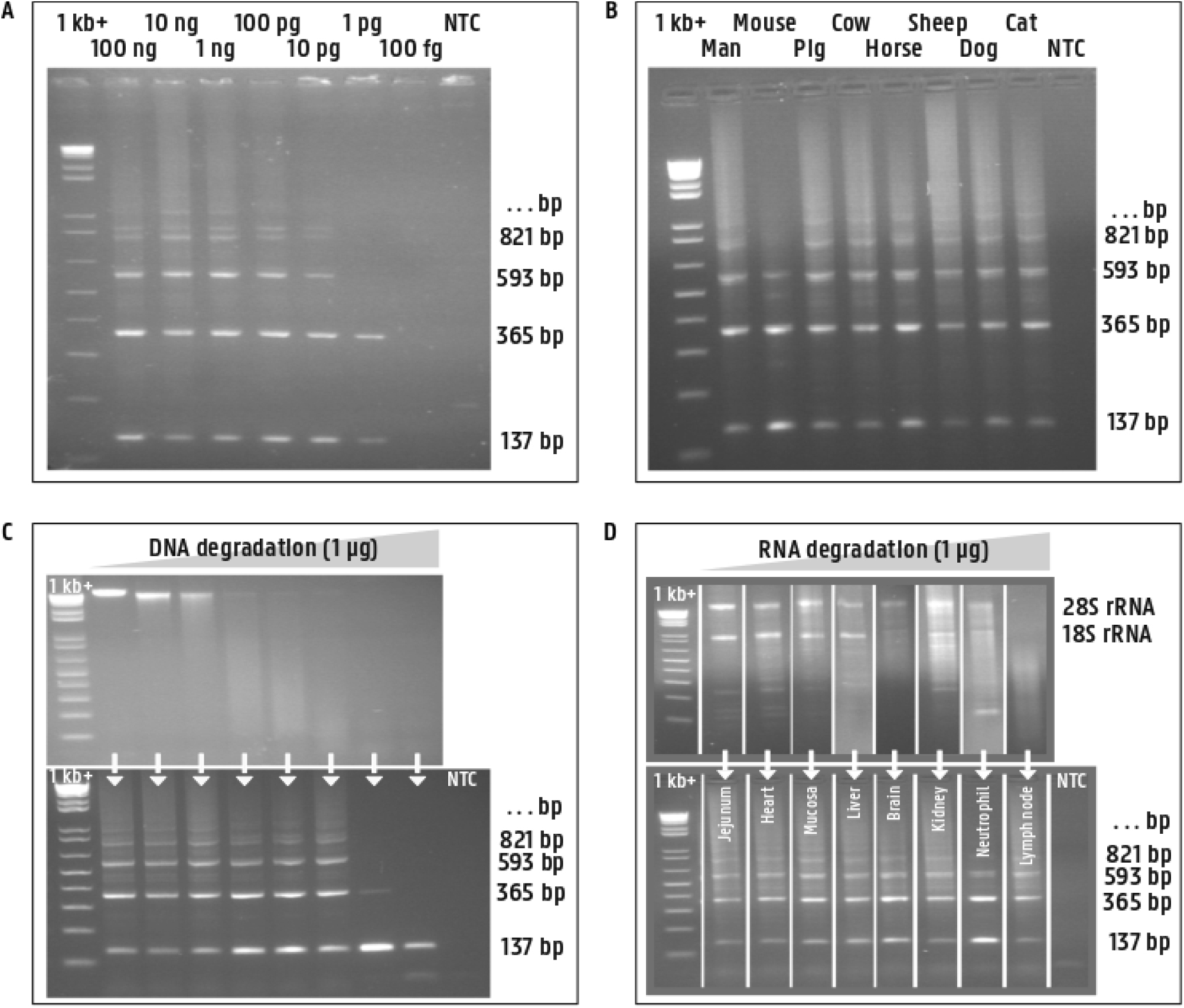
Agarose gels showing amplicons generated with the UBC integrity assay on (A) a 1/10 serial dilution series of porcine DNA, (B) 100 ng DNA isolated from 8 mammal species, (C) 10 ng porcine DNA gradually degraded with DNase (1 μg gradually degraded DNA is analysed in the upper part, the assay on the corresponding samples in the lower part) and (D) cDNA reverse transcribed from RNA from 8 different tissues with different quality (1 μg RNA is analysed in the upper part, the assay on cDNA from the corresponding RNA in the lower part).

In order to evaluate if decreasing quality of nucleic acids results in a decrease of both the number and the concentration of the amplicons starting with the longer fragments, the assay was performed on high quality DNA gradually degraded with DNase (Figure 2.C) and on cDNA reverse transcribed from RNA with different degradation levels (Figure 2.D). Surprisingly, for both degraded DNA and RNA, high amplificability was observed, showing that even heavily degraded nucleic acids will still be useful for a lot of applications.

The assay was found to be robust since it was also successfully applied to evaluate DNA from equine vaginal swabs, DNA from faeces from an unknown origin, circulating DNA from equine plasma (cfDNA), contaminating DNA in porcine RNA samples (minus RT control PCR) and cDNA from bovine embryos, and performed by different scientists with different polymerases on different PCR thermal cyclers (data will be published elsewhere).

### Validation of the UBC bisulfite integrity assay

DNA was isolated from 8 biological replicates of bovine blood neutrophils. The UBC integrity assay showed that the DNA was of high quality because all desired amplicons were generated (Figure 3.A). After bisulfite treatment, known to damage DNA severely, these samples were used to validate the UBC bisulfite integrity assay. Gradient PCR showed that a Ta range of 52-56°C gave the best results (54°C was chosen as optimal Ta; Additional file 5.B). The optimized assay was performed on the 8 biological replicates. Surprisingly, not only the 2 lower fragments of 105 and 333 bp were amplified, but also longer amplicons, although less intense (Figure 3.B). This finding suggest that amplicons longer than the recommended 300 bp could be used for bisulfite sequencing. To verify this, 2 bovine specific assays for bisulfite sequencing, amplifying 442 bp of MPO and 462 bp of SOD2, were performed on the bisulfite treated DNA of the 8 biological replicates. Agarose gel analysis showed that both amplicons were successfully amplified on all samples (Figure 3.C-D).

**Figure 3.**
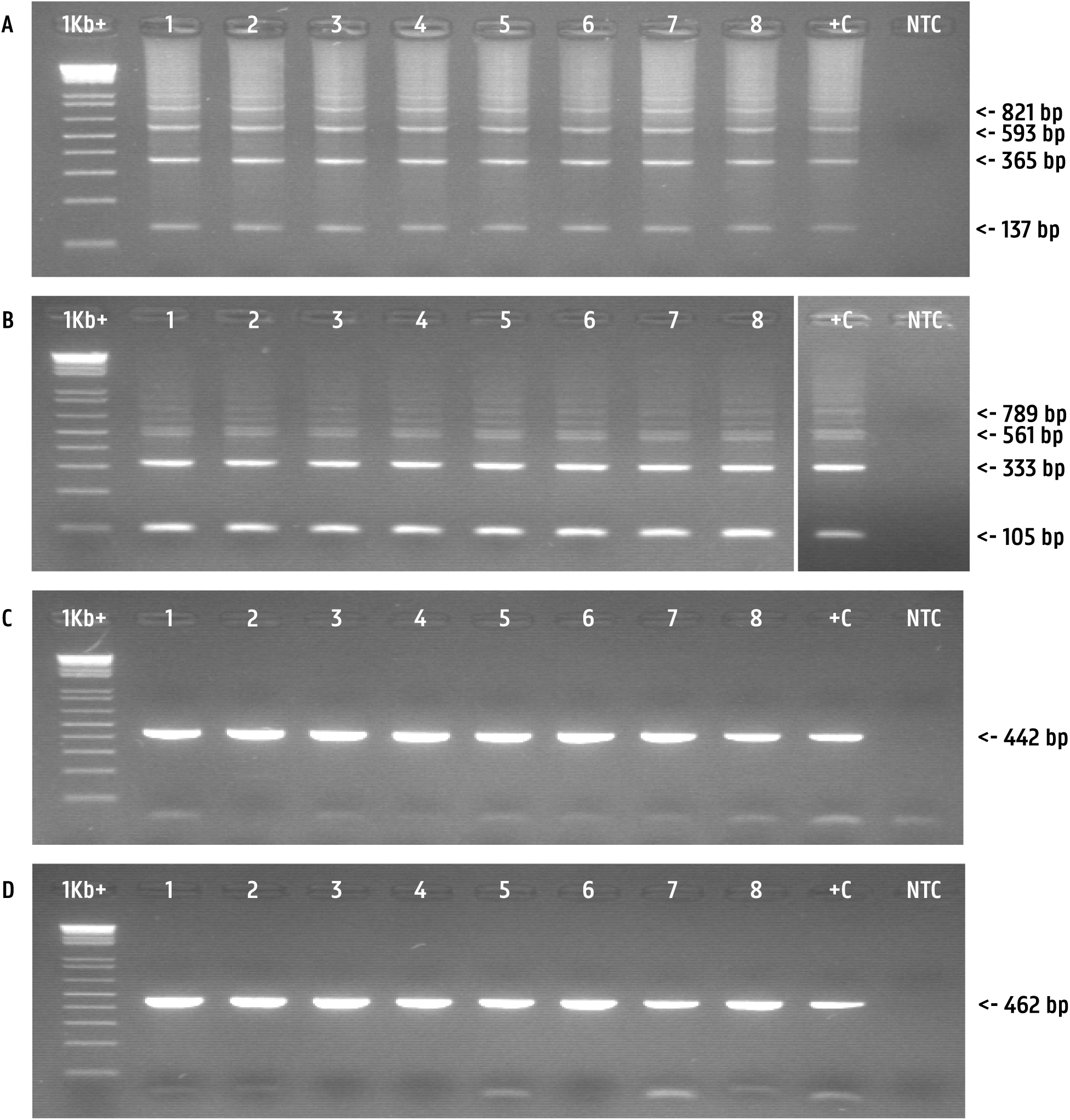
Agarose gels showing amplicons generated with (A) the UBC integrity assay on native DNA isolated from 8 biological replicates of bovine blood neutrophils, and with (B) the UBC bisulfite integrity assay, (C) the *Btau*MPO assay and (D) the *Btau*SOD assay on the corresponding DNA samples after bisulfite treatment.

## CONCLUSIONS

We conclude that UBC gene features are extremely suited to develop single, cheap and multi-use PCR-based assays to estimate the presence, integrity and amplificability of DNA (isolated from a variety of sample types/qualities from different species) and cDNA (reverse transcribed from RNA isolated from any tissue type, reflecting the RNA integrity and taking into account the efficacy of the reverse transcription). In this paper we describe a single assay useful to investigate native nucleic acids from all tissues of the most investigated mammal species. Surprisingly, the PCR amplificability was a lot higher than expected from electrophoresis-based integrity assay results, what could be of great help for validating degraded samples for (non-)quantification applications. This assay can also be used to check the level of DNA contamination in RNA samples (minus RT control PCR) or to check if an unknown sample contains useful mammalian DNA for further investigation (forensics, species identification). We strongly believe that this assay will prove its usefulness and will contribute to the improvement of quality control strategies. Laboratories that need to check PCR amplificability might directly benefit from this single integrity assay (instead of using a battery of more specific integrity assays) or apply the same strategy to design similar multiuse assays, as we did for estimating the integrity of bisulfite treated DNA from the most investigated mammal species.

## ACKNOWLEDGEMENTS

We wish to thank Laice Angel Royeras, Caro Rogiers, Dominique Vander Donckt, Linda Impe and Ruben Van Gansbeke for excellent technical assistance.

